# Ten-year retrospective analysis of *Acinetobacter baumannii* clinical isolates reveals a proportionately large, non-nosocomial, multidrug-resistant endemic reservoir

**DOI:** 10.1101/576868

**Authors:** Juan J Calix, Jason P Burnham, Mario F Feldman

**Affiliations:** Division of Infectious Diseases, Department of Medicine, Washington University School of Medicine, St. Louis, MO; Department of Molecular Microbiology, Washington University School of Medicine, St. Louis, MO

**Keywords:** *Acinetobacter baumannii*, multidrug resistance, hospital-acquired infections

## Abstract

**Background:** *Acinetobacter baumannii* (*Ab*) is a global health threat notorious for causing hospital-acquired (HA) infections, though many *Ab* infections are community-acquired (CA). Investigations describing contemporaneous, clinically-relevant CA and HA *Ab* populations, are lacking.

**Methods:** We conducted a retrospective ecological analysis of 2042 *Ab* clinical isolates identified from 2007 to 2017 in the BJC HealthCare System (BJC), a multi-hospital system located in and around the greater metropolitan area in St. Louis, Missouri. We described basic clinical characteristics and antibiotic susceptibility rates of CA and HA *Ab* isolates in comparative and longitudinal analyses.

**Results:** 62.1% of all *Ab* isolates were CA, i.e., isolated in ambulatory settings or <48 hours following hospital admission. Though HA isolates initially predominated in the largest BJC hospital, implementation of infection control efforts resulted in a disproportionate reduction in annual HA isolate occurrence. This revealed a stable, baseline occurrence of CA isolates. In all other hospitals, the annual proportion of isolates that were CA averaged 78.7% (95%CI=74.5-83.0). 42.9% and 30.4% of total CA isolates were from skin and soft tissue/musculoskeletal (SST/MSK) and urinary sources, respectively, while HA isolates were primarily respiratory (55.6%). Rates of carbapenem resistance, a surrogate for multidrug resistant (MDR) phenotypes, were higher among respiratory and HA cases (∼60%) compared to contemporaneous non-respiratory CA counterparts (∼40%).

**Conclusions:** MDR *Ab* reservoirs associated with SST/MSK and urinary niches persist outside of hospital environments in a large U.S. healthcare system, even after the implementation of effective hospital infection control measures.

**Summary:** We compared hospital-acquired and community-acquired *Acinetobacter baumannii* in a large U.S. healthcare system through a ten-year retrospective ecological analysis. Community-acquired isolates composed over 60% of total *A. baumannii* isolates, were primarily from non-respiratory sources and exhibited carbapenem resistance rates of 35-40%.

## Introduction

The gram-negative bacterium *Acinetobacter baumannii* (*Ab*) can survive in multiple host and abiotic environments and exhibits a propensity to acquire resistance to most antibiotics, including carbapenems [1, 2]. In response to the global impact of multidrug resistant (MDR) *Ab* infections, the World Health Organization and U.S. Centers for Disease Control and Prevention have recognized *Ab* as an urgent threat requiring the development of novel interventions [3, 4]. However, the epidemiology and pathogenesis of clinically-relevant *Ab* remain incompletely characterized.

*Ab* is widely regarded as an opportunistic pathogen that rarely causes community-acquired (CA) infections, but instead causes hospital-acquired (HA) infections, namely nosocomial pneumonia and bacteremia, in critically ill or immunocompromised patients [2, 5-7]. Thus, *Ab*-related research has almost exclusively focused on hospital-associated bacterial populations [8-14]. However, recent studies suggest that *Ab* isolates are routinely acquired in outpatient settings [15-18]. Therefore, research biased towards HA cases may fail to describe the full spectrum of clinically-relevant *Ab* reservoirs. Specifically, there is a paucity of investigations comparing contemporaneous *Ab* populations with differing epidemiological traits, such as CA versus HA isolates, or isolates from large academic versus community hospitals. Defining these potentially divergent *Ab* populations is especially important for accurately gauging the impact of interventions designed against HA infections.

Here, we characterized different *Ab* populations through a retrospective longitudinal analysis of *Ab* isolates identified over ten years in a large, multi-hospital system in St. Louis, Missouri. Notably, an effective campaign against nosocomial *Ab* infections, which included the 2012 relocation of an ICU implicated in multiple *Ab* outbreaks, occurred during this period, allowing us to observe the impact of these interventions on different *Ab* populations. In this study, we did not investigate clinical outcomes or patient-specific risk factors, instead focusing on comparing isolate-associated clinical features to better understand the ecology of clinically-relevant *Ab* populations. Using this approach, we identified clinical features distinguishing *Ab* populations and confirmed the emerging impact of CA *Ab*.

## Methods

### Study Location and Period

Following approval from our local Institutional Review Board, we performed a retrospective analysis of isolates identified in the BJC HealthCare System (BJC) from January 2007 to September 2017. BJC is a large integrated inpatient and outpatient healthcare system in and around the Greater Metropolitan Area of St. Louis, Missouri, USA. It includes nine community hospitals, an academic pediatric hospital and a 1250-bed academic adult medical center (Barnes-Jewish Hospital, BJH), which all combine for a total of over 3200 inpatient beds and >140 000 admissions annually (**Table S1**). For longitudinal analyses we used data only from 2007 to 2016, given that 2017 data was limited to January through August, at time of analysis.

### Isolate Identification and Definitions

Using the BJC Clinical Data Repository (CDR), which is maintained by the BJC Center for Clinical Excellence, we identified all instances in which *Acinetobacter* was isolated during the course of regular medical care from adult patients age ≥18 years. Surveillance cultures obtained during suspected nosocomial outbreaks were excluded. Only isolates from the first isolation event per patient (“index culture”) was eligible for inclusion. Isolates were identified using either automated biochemical methods or matrix-assisted laser desorption/ionization and time of flight spectroscopy. The number of *Acinetobacter* index cultures and their microbial identification are listed in **Table S2**. Only cases identified as “*Acinetobacter baumannii*” (n=990) or “*Acinetobacter calcoaceticus-baumannii* complex” (n=1052) were combined for the current analysis. Basic patient demographic information, isolate tissue source, hospital day of index culture (if applicable), and antibiotic susceptibility data for each isolate was obtained from the BJC CDR and by review of electronic charts, as needed. Isolates were classified into one of five anatomical categories according to isolate tissue source: “respiratory”, “skin and soft tissue/musculoskeletal” (SST/MSK), “urinary”, “endovascular”, or “other.” They were defined as “hospital-acquired” (HA) if index culture was performed ≥48 hours after hospital admission and prior to discharge, while all other isolates were defined as “community-acquired” (CA). Isolates were also classified as “multi-isolate” if >1 co-isolated microbial species was reported in index culture, or “sole isolate” if only a single *Acinetobacter* isolate was reported in the index culture.

### Antibiotic susceptibility reporting

Antibiotic susceptibility testing was performed using the Vitek 2 system or Kirby-Bauer disk diffusion on Mueller-Hinton Agar, and interpreted per CLSI guidelines [19]. Due to temporal and geographical variation in susceptibility testing practices, antibiotic susceptibility profiles were incomplete for many isolates. Isolates lacking susceptibility reporting for an antibiotic were excluded from respective susceptibility-associated analyses. Non-susceptible isolates (i.e., isolates reported as “resistant” or “intermediate”) were classified as “resistant” for analyses. Lastly, if an isolate was non-susceptible to any for the following antibiotics in a class, it was labeled “resistant” for that class: imipenem or meropenem for “carbapenems”; ciprofloxacin or levofloxacin for “fluoroquinolones”; piperacillin-tazobactam or ticarcillin-clavulanic acid for “antipseudomonal penicillins plus β-lactamase inhibitor”; and tetracycline or doxycycline for “tetracyclines” (**Table S3**).

### Statistical Methods

All analyses were performed with SPSS v25 (IBM, USA). Chi-squared test or independent t-test was performed for comparing categorical or continuous variables, respectively. *P* values <0.05 were considered statistically significant.

### Results

#### Isolates from different hospitals exhibit separate longitudinal trends

Of the 2042 eligible *Ab* isolates obtained in BJC hospitals from January 2007 through September of 2017, 48.3% (n=987) were obtained at BJH (Table S1). The remaining isolates were identified in various smaller hospitals (herein, referred to as “non-BJH” hospitals). As seen in **Figure 1A**, annual *Ab* occurrence at BJH increased from 2007 to 2009, and steadily decreased over the remainder of the study period. In contrast, annual occurrence of non-BJH isolates was relatively constant. Given this differential pattern, we grouped isolates as “BJH” and “non-BJH” in our longitudinal analyses.

**Figure 1.**
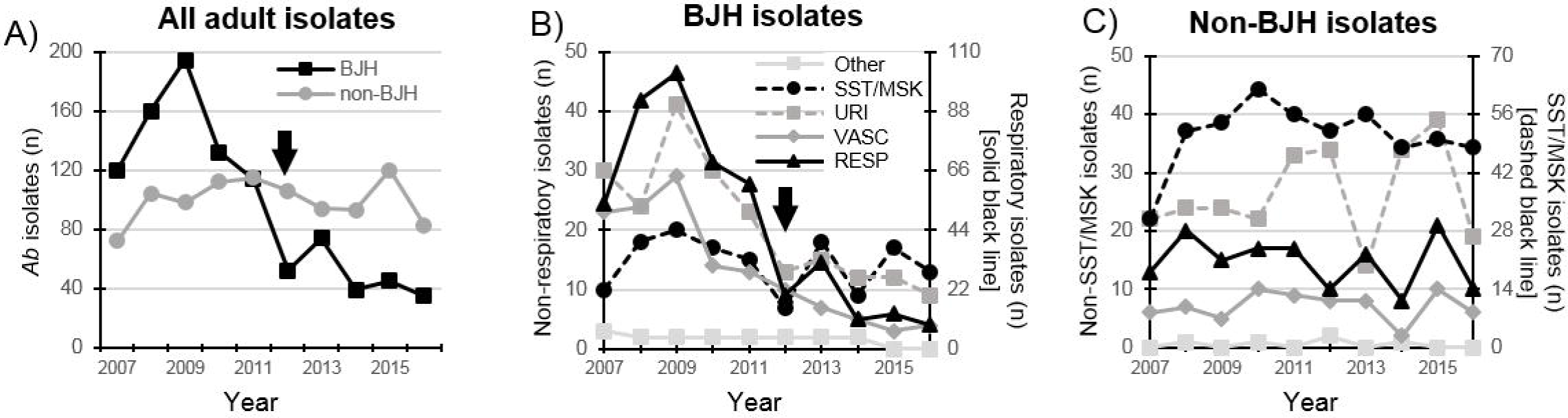
Annual occurrence of adult Ab isolates, BJC 2007-2016. Panel A depicts annual BJH (black) and non-BJH (gray) Ab isolates. Also depicted are the annual amounts of BJH (panel B) and non-BJH (panel C) adult Ab isolates obtained from each anatomic source. The legend in panel B applies to both panels B and C. Values for BJH respiratory (solid black line) and non-BJH skin and soft tissue/musculoskeletal (SST/MSK, dashed black line) isolates are on the right y-axis in panel B and C, respectively. All other values are on the left y-axis. Arrows depict the year during which a BJH ICU implicated in multiple nosocomial Ab outbreaks was relocated (see text). RESP, respiratory; URI, urinary; VASC, endovascular.

#### Adult *Ab* isolates were derived from various anatomical sources

Contrary to the prevalent notion that *Ab* is predominantly a respiratory pathogen [2, 5], 692 isolates (33.9% of all adult isolates) were from skin and soft tissue/musculoskeletal (SST/MSK) sources, while 626 (30.7%), 487 (23.8%), 214 (10.5%), and 23 (1.1%) isolates were from respiratory, urinary, endovascular, and “other” sources, respectively (Table 1). Proportion of “sole isolate” cases, where *Ab* was the only isolate in index culture, differed across anatomic sources (*p*<0.001), with endovascular and SST/MSK compartments having the highest and lowest proportions (83.2% and 51.5%, respectively). In longitudinal analysis, annual BJH respiratory, urinary, and endovascular isolates peaked in 2009 and subsequently decreased ∼70% by 2016 (**Figure 1B**), with the largest year-over-year decrease happening in 2012 (**Figure 1B, arrow**). In contrast, annual BJH SST/MSK isolates and non-BJH isolates from all sources remained relatively stable (**Figures 1B and 1C**). Thus, isolates from different anatomic sources and different hospitals exhibited varying epidemiologic features.

**Table 1.**
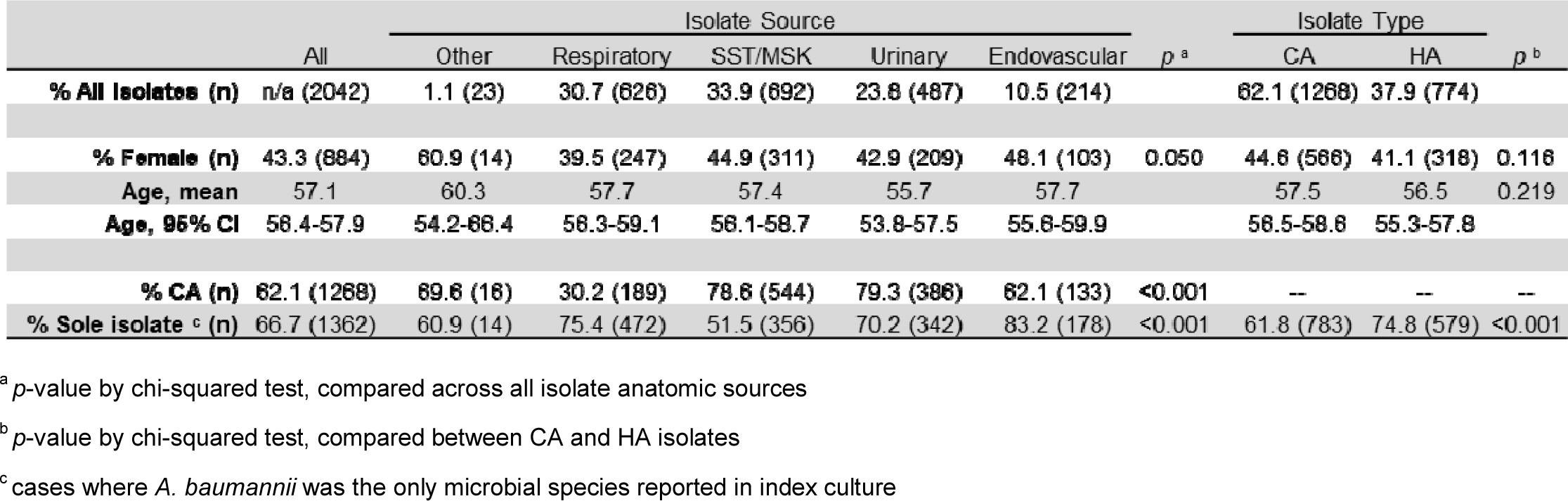
General clinical characteristics of all adult *A. baumannii* isolates, BJC 2007-2017.

|  | All | Isolate Source |  |  |  |  | *p*<sup>a</sup> | Isolate Type |  | *p*<sup>b</sup> |
| --- | --- | --- | --- | --- | --- | --- | --- | --- | --- | --- |
|  |  | Other | Respiratory | SST/MSK | Urinary | Endovascular |  | CA | HA |  |
| **% All Isolates (n)** | n/a (2042) | 1.1 (23) | 30.7 (626) | 33.9 (692) | 23.8 (487) | 10.5 (214) |  | 62.1 (1268) | 37.9 (774) |  |
| **% Female (n)** | 43.3 (884) | 60.9 (14) | 39.5 (247) | 44.9 (311) | 42.9 (209) | 48.1 (103) | 0.050 | 44.6 (566) | 41.1 (318) | 0.116 |
| **Age, mean** | 57.1 | 60.3 | 57.7 | 57.4 | 55.7 | 57.7 |  | 57.5 | 56.5 | 0.219 |
| **Age, 95% CI** | 56.4-57.9 | 54.2-66.4 | 56.3-59.1 | 56.1-58.7 | 53.8-57.5 | 55.6-59.9 |  | 56.5-58.8 | 55.3-57.8 |  |
| **% CA (n)** | 62.1 (1268) | 69.6 (16) | 30.2 (189) | 78.6 (544) | 79.3 (386) | 62.1 (133) | <0.001 | -- | -- | -- |
| **% Sole isolate<sup>c</sup> (n)** | 66.7 (1362) | 60.9 (14) | 75.4 (472) | 51.5 (356) | 70.2 (342) | 83.2 (178) | <0.001 | 61.8 (783) | 74.8 (579) | <0.001 |

<sup>a</sup> *p*-value by chi-squared test, compared across all isolate anatomic sources
<sup>b</sup> *p*-value by chi-squared test, compared between CA and HA isolates
<sup>c</sup> cases where *A. baumannii* was the only microbial species reported in index culture

#### HA and CA isolates exhibited divergent epidemiology

Of all adult *Ab* isolates, 37.9% (n=774) were HA and 62.1% (n=1268) were CA (**Table 1**). The percent of all adult isolates that were CA (“CA ratio”) increased over the study period (**Figure 2A**) and varied among hospitals (**Table S1**). Notably, the decline in annual BJH *Ab* isolates over the study period (**Figure 1A**) was largely due to a >10-fold decrease in annual HA isolates from 2009 to 2016 (**Figure 2B**). Though annual BJH CA isolates also exhibited a ∼3-fold decrease from 2009 levels, they remained relatively stable after 2012. The decline of HA *Ab* occurrence resulted in the BJH annual CA ratio increasing from 39.2% to 74.3% over the study period (**Figure 2B and Table S1**). In contrast, annual CA ratios among non-BJH isolates remained largely unchanged (mean= 78.7%; 95%CI=74.5-83.0) (**Figure 2C**).

**Figure 2.**
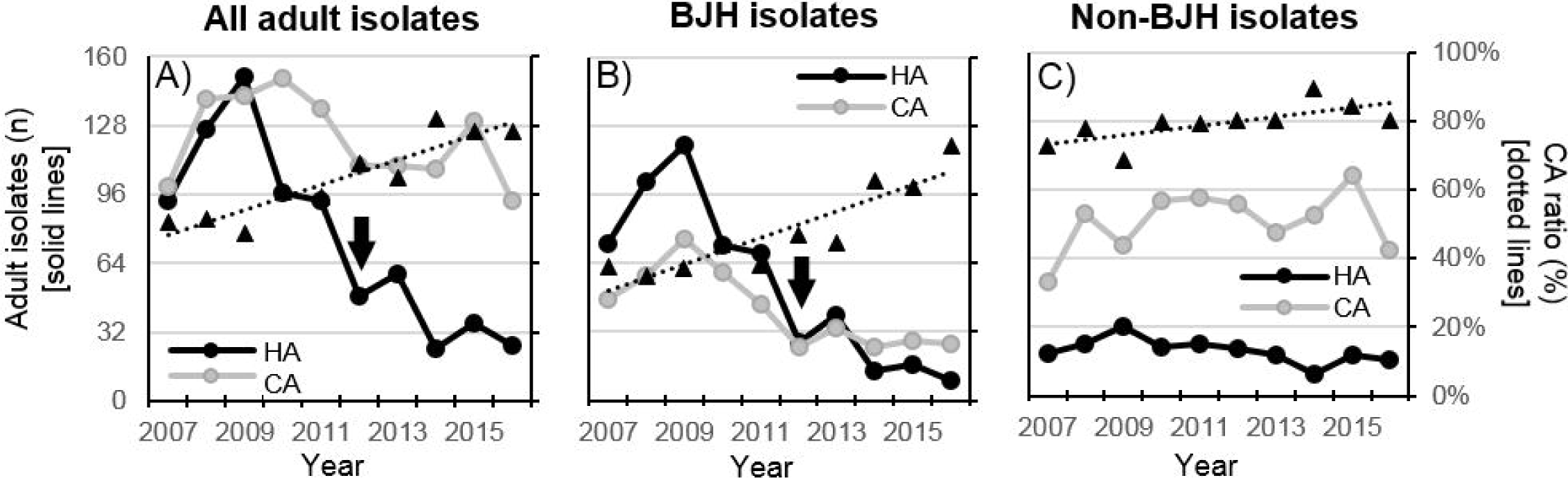
Annual amounts of community-acquired (CA, gray) and hospital-acquired (HA, black) isolates among all (panel A), BJH (panel B) and non-BJH (panel C) adult isolates. In all panels, black triangles depict annual percent of isolates that are CA (“CA ratio”) and dotted lines are a best-fit trend lines for CA rates (values on right y-axis). Y-axis values are conserved across panels. Arrows depict the year during which a BJH ICU implicated in multiple nosocomial Ab outbreaks was relocated (see text).

The comparable selective decline in annual BJH HA and respiratory *Ab* isolates (**Figure 1B**), suggested a link between these epidemiologic compartments. Indeed, 56.5% of total HA isolates were from respiratory sources, followed by SST/MSK (19.1%), urinary (13.0%), endovascular (10.5%) and “other” (0.9%). In contrast, CA isolates were primarily SST/MSK (42.9%) and urinary (30.4%), with only 14.9%, 10.5% and 1.3% isolated from respiratory, endovascular and “other” sources, respectively. Similarly, 79.3%, 78.6%, and 62.1% of total urinary, SST/MSK, and endovascular isolates, respectively, were CA, compared to only 30.2% of respiratory isolates (**Figure 3**). CA ratios were higher among non-BJH isolates in each anatomic source category, but the association between anatomic source and CA ratio was conserved in both BJH and non-BJH isolates (**Figure 3**).

**Figure 3.**
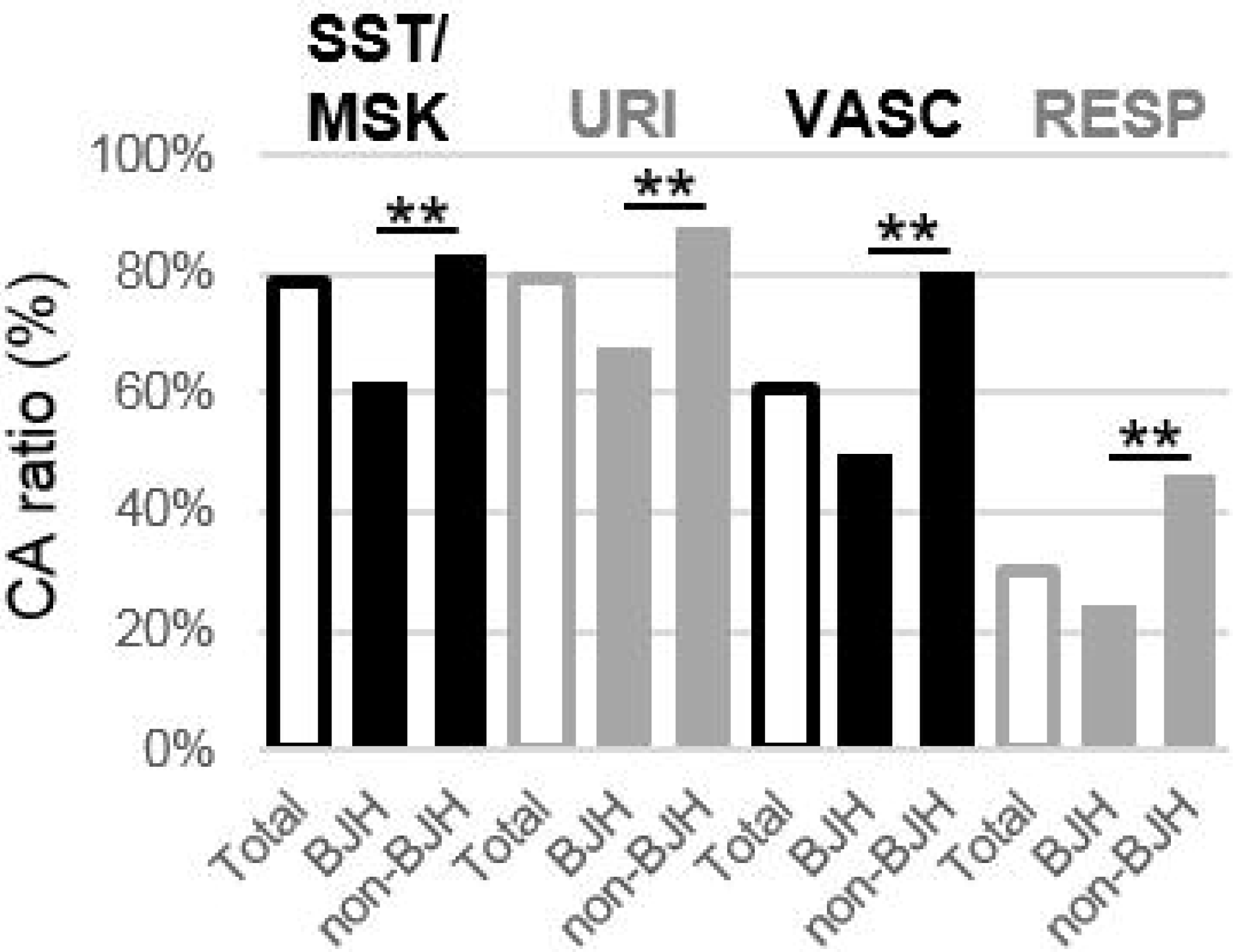
Percent of total, BJH and non-BJH adult isolates that were CA (“CA ratio”). Isolates are grouped by anatomical source: skin and soft tissue/musculoskeletal (SST/MSK), urinary (URI), endovascular (VASC) or respiratory (RESP). White bars correspond to “total” adult isolates for each group. **, p<0.005 by chi-squared test.

As seen in **Figure 4**, annual CA ratios were relatively stable over time for non-BJH SST/MSK, urinary and respiratory isolates, with mean CA ratios of 83.3% (95%CI=78.7-87.8), 87.5% (95%CI=82.4-92.6) and 46.8% (95%CI=41.2-52.4), respectively. Though annual CA ratios for BJH SST/MSK isolates were also relatively unchanged (mean=60.4%, 95%CI=41.2-52.4), CA ratios changed over time for other BJH isolate types. From 2007-2011, annual CA ratios for BJH respiratory and urinary isolates averaged 23.5% (95%CI=20.6-26.4) and 61.8% (95%CI=57.1-66.5), respectively. Annual CA rates varied among BJH respiratory isolates from 2012-2016, averaging 35.6% (95%CI=8.8-62.5). Contemporaneously, there were nine to eleven annual CA isolates from 2012-2016, while annual HA urinary isolates declined to zero. This resulted in BJH urinary isolate CA ratios progressively increasing to 100% in 2016 (**Figure 4**). Both BJH and non-BJH endovascular isolates exhibited gradual increases in CA ratios that began prior to 2012 (**Figure 4**), with CA ratios among all endovascular isolates increasing from 44.8% (13 of 29 isolates) in 2007 to 100% in 2014 and 2015 (n=7 and 13, respectively) and 60% in 2016 (6 of 10 isolates). In summary, the BJC clinically-relevant *Ab* population was predominated by CA isolates principally from urinary and SST/MSK sources, and their occurrence was largely independent of HA isolates, which were principally from respiratory sources. Furthermore, a decrease in annual HA isolates overall (**Figure 2A**), coincided with increases in CA ratios among endovascular isolates (**Figure 4**).

**Figure 4.**
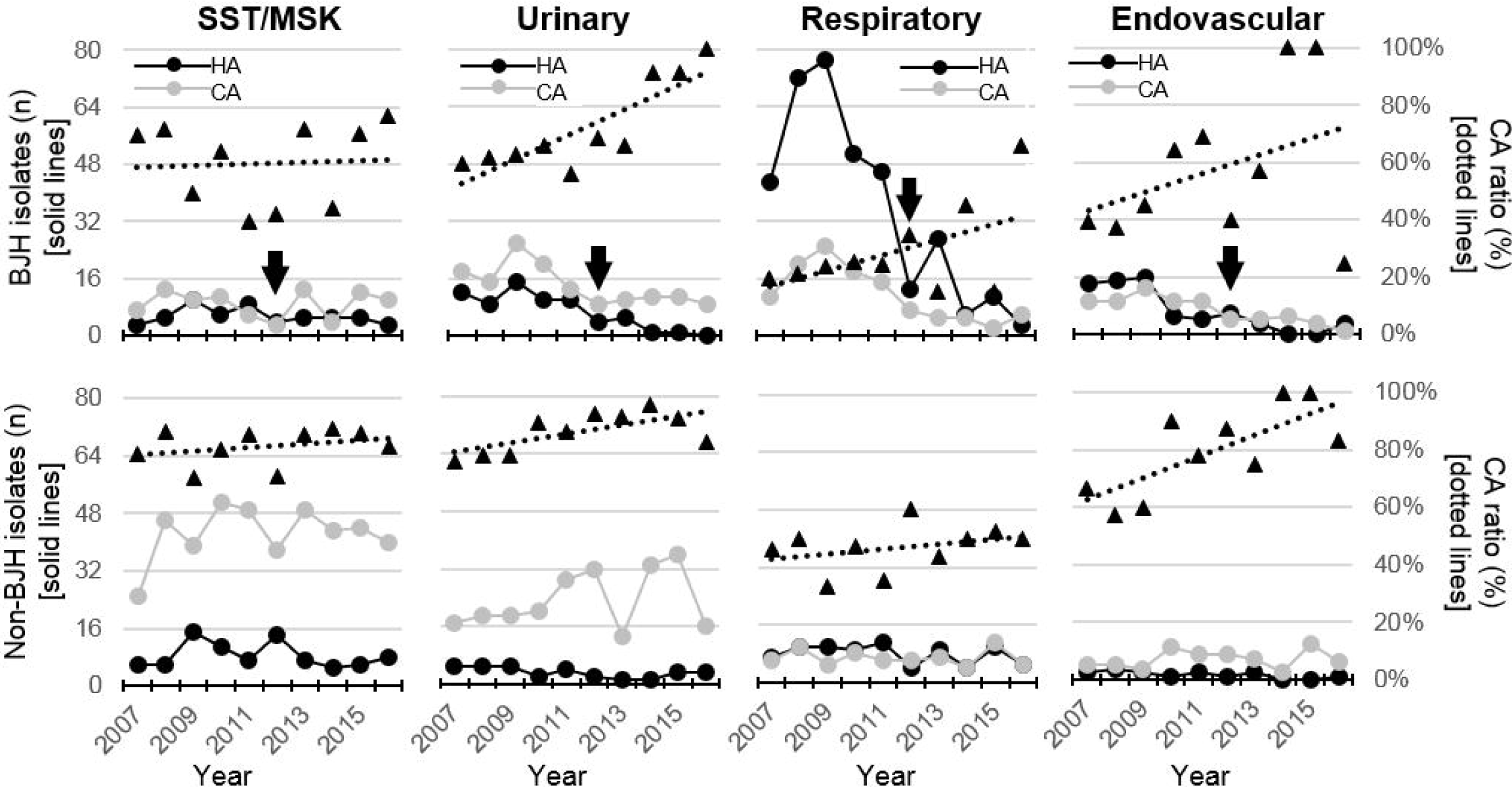
Prevalence of community-acquired isolates differs by Ab isolate source. Annual amounts of hospital-acquired (HA, black lines) and community-acquired (CA, gray lines) Ab isolates in BJH (top row) and non-BJH hospitals (bottom row). Columns correspond to isolates obtained from each anatomic source. Y-axis scale is maintained throughout graphs in a row. In all panels, triangles depict annual percent of isolates that are CA (“CA ratio,” values on right y-axis), and dotted lines are a best-fit trend lines for CA ratio values. Arrows depict the year during which a BJH ICU implicated in multiple nosocomial Ab outbreaks was relocated (see text). Isolates from “other” sources are omitted for clarity. SST/MSK, skin and soft tissue/musculoskeletal.

#### High prevalence of antibiotic resistance among adult *Ab* isolates

As shown in **Table S3**, adult *Ab* isolates exhibited high rates of antibiotic resistance, with rates ranging from 27.5% for gentamicin to 90.5% for ceftriaxone. Antibiotic resistance was associated with multiple clinical characteristics, including being “sole isolate” in index culture and older patient age. Resistance rates also differed between HA and CA isolates and among isolates from different anatomic sources. However, with the exception of ampicillin-sulbactam, there were less than two-fold differences between the high resistance rates exhibited by HA, respiratory and endovascular isolates, and the lower rates among CA, urinary and SST/MSK isolates (**Table S3**). Therefore, adult *Ab* isolates in all compartments exhibited elevated antibiotic resistance rates.

#### Rate of carbapenem resistance, a marker for *Ab* MDR phenotypes, varied according to epidemiologic compartment

Since *Ab* susceptibility testing practices varied in BJC during this period, we could not reliably determine whether an isolate met established MDR definitions [20], i.e., non-susceptibility to ≥1 agent in ≥3 antimicrobial categories (**Table S3, first row**). However, all 867 adult carbapenem resistant *Ab* (CRAb) isolates were resistant to at least two other antibiotic classes, independent of ceftriaxone (data not shown). Thus, as previously observed in other *Acinetobacter* populations [21], carbapenem resistance was a marker of the MDR phenotype. Annual rate of carbapenem resistance (“CRAb-rate”) ranged from 34.2% in 2012 to 58.9% in 2009 among total *Ab* isolates. Annual CRAb-rates differed between total HA and CA isolates, averaging 38.1% (95%CI=32.7-43.5) and 56.3% (95%CI=49.0-63.5), respectively (**Figure 5A**). BJH isolate CRAb-rates markedly changed in 2012, with an average of 58.3% (95%CI=51.6-65.1) from 2007-2011, and 36.6% (95%CI=32.2-41.0) from 2012-2016 (**Figure 5B**). In contrast, CRAb-rates among non-BJH adult isolate were stable throughout the study period at 39.3% (95%CI=34.2-44.5) (**Figure 5C**). In summary, HA isolates had stably higher CRAb-rates than CA isolates, and total *Ab* CRAb-rates changed over time, according to the prevalence of HA and CA isolates in the population.

**Figure 5.**
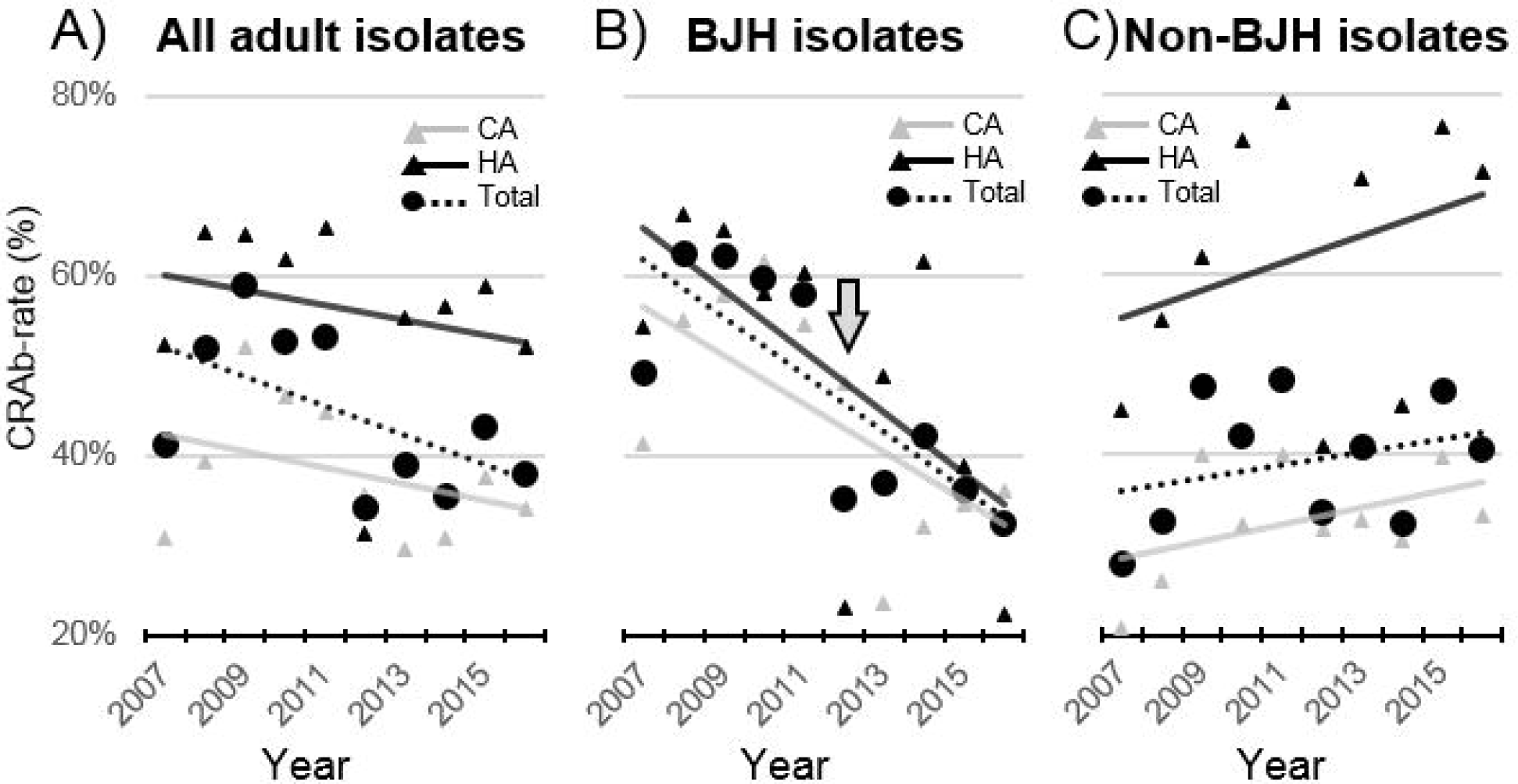
Rates of carbapenem resistance (CRAb-rate) among all (panel A), BJH (panel B) and non-BJH (panel C) adult *Ab* isolates, BJC 2007-2016. Annual CRAb-rates among all (black circles, dashed line), hospital-acquired (black triangles and solid line), and community-acquired (gray diamonds and solid line) *Ab* isolates. Arrow depicts the year during which a BJH ICU implicated in multiple nosocomial *Ab* outbreaks was relocated (see text).

CRAb-rates were comparable among HA isolates from different anatomic sources (**Figure 6A, black bars**). Furthermore, CRAb-rates were indistinguishable between CA and HA respiratory isolates (61.2% and 55.8%, respectively, *p*=0.22). In contrast, CRAb-rates were lower in CA versus HA isolates from SST/MSK (36.7% and 63.4%, respectively), urinary (30.8% and 61.1%), and endovascular (41.2% and 65.3%) sources (*p*<0.001 for all comparisons) (**Figure 6A**). The dissimilar CRAb-rates among non-respiratory CA and HA populations were present throughout the period (**Figure S1**), and observed among both BJH and non-BJH isolates (**Figure S2**). Thus, there were two populations according to diverging CRAb-rates (**Figure 6A**) – “highly resistant” populations with CRAb-rates >55%, i.e., all HA isolates and CA respiratory isolates; and “intermediately resistant” populations with CRAb-rates between 20-50%, i.e., non-respiratory, CA isolates.

**Figure 6.**
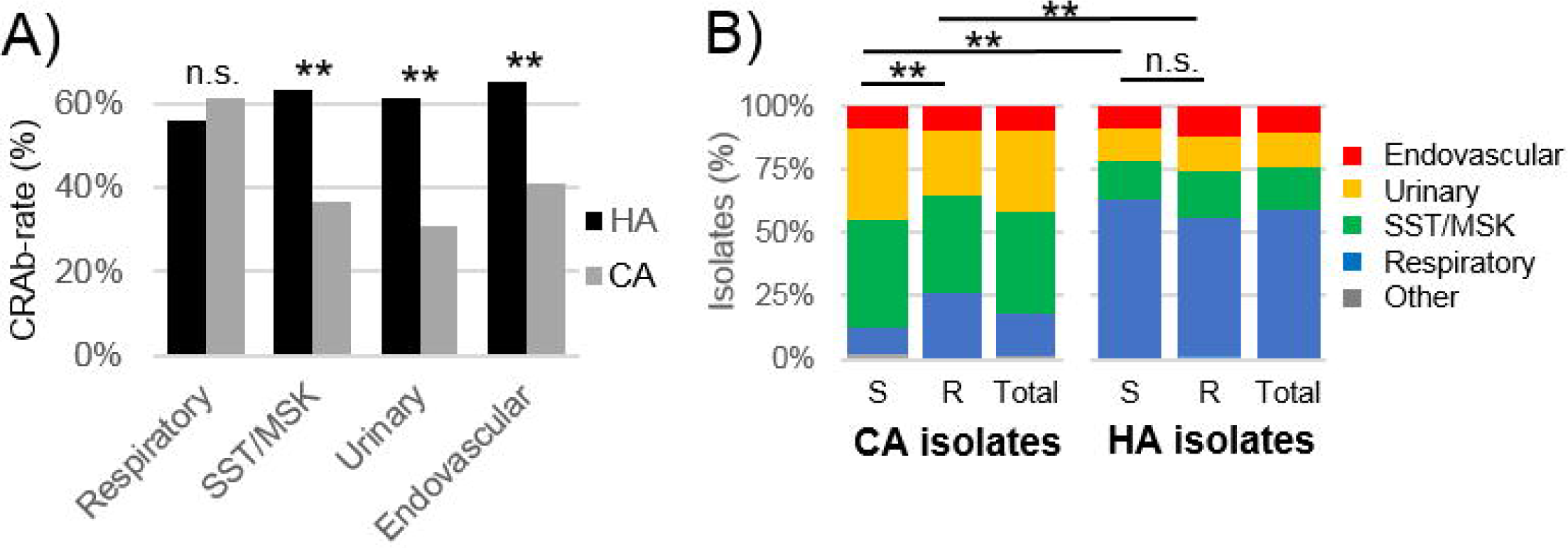
Rates of carbapenem resistance (CRAb-rate) differ according to *Ab* isolate type. A) CRAb-rates among isolates from each anatomic source, grouped by hospital-acquired (HA, black) and community-acquired (CA, gray) cases. CRAb-rate was compared between HA and CA isolates by chi-squared test. B) Proportion of carbapenem-susceptible (S), -resistant (R) or total adult *Ab* isolates from each anatomic source. Isolates were grouped into HA and CA. The proportion of isolates from each source was compared between compartments by chi-squared test. SST/MSK, skin and soft tissue/musculoskeletal; n.s., not significant; **, *p*<0.005.

When comparing the proportions of isolates from each anatomic source, there was no difference between CRAb and carbapenem-susceptible HA isolates (*p*=0.77) (**Figure 6B**). Though the proportions differed between susceptible and resistant CA isolates (*p*<0.005), this difference was minimal compared to the dissimilarities between HA and CA isolates, independent of carbapenem susceptibility (**Figure 6B**). Respiratory isolates composed 55.0% of HA CRAb isolates but only 25.8% of total CA CRAb isolates. Conversely, SST/MSK and urinary isolates composed 40.8% and 31.9%, respectively, of the CA CRAb isolate reservoir, while composing only 18.5% and 13.7%, respectively, of the HA CRAb reservoir. Endovascular isolates composed ∼10% of all compartments (**Figure 6B**). In summary, CA CRAb/MDR isolates were principally from urinary and SST/MSK sources, while HA CRAb/MDR isolates were principally from respiratory sources.

### Discussion

Antibiotic-sparing strategies against MDR *Ab* must target factors facilitating bacterial survival in pertinent reservoirs from where *Ab* infects at-risk hosts. To better characterize the ecology of *Ab* reservoirs, we retrospectively analyzed 2042 temporally- and geographically-associated *Ab* clinical isolates. In contrast to the widely accepted notion that *Ab* is primarily a HA pathogen, we found that 60-80% of *Ab* isolates were CA. This high CA ratio may result from the inclusion of multiple regional community hospitals, resulting in a more comprehensive survey of local *Ab* reservoirs. Indeed, if we had surveyed only our large academic center, BJH, the CA ratio would have been <45% (**Table S1**). Another possible explanation for a high CA ratio may be that multiple CA isolates were obtained through unaccounted healthcare exposures, such as recent hospitalizations or long-term acute care facilities [22]. A limitation to this study is that we could not identify patients who were transferred from non-BJC facilities or who had other risk factors that would classify their cases as “healthcare-associated” (HCA) [23]. However, multiple similar studies have reported that 25-65% of *Ab* clinical isolates are likely acquired in the community [15-18], supporting that a substantial portion of clinically-relevant *Ab* reside in outpatient settings.

Further affirming the existence of an endemic *Ab* community reservoir, the occurrence of CA isolates persisted even after the near eradication of BJH HA *Ab* cases. Multiple HA isolates identified in 2007-2012 were from patients in a BJH ICU implicated in several MDR *Ab* nosocomial outbreaks starting in late 2007 and ending in August 2011 (unpublished findings). The 10-fold decrease of annual BJH HA isolates likely resulted from physical relocation of the suspect ICU ward in 2012 and other aggressive hospital-wide infection control measures. We suspect the accompanying 3-fold decrease in CA isolates was secondary to a reduction of unaccounted HCA cases. While annual HA respiratory, urinary and endovascular isolates decreased to near-zero levels after 2012, there was a steady annual occurrence of CA isolates with epidemiologic features similar to CA isolates from non-BJH hospitals (i.e., intermediately carbapenem resistant isolates from urinary and SST/MSK sources). Thus, the selective decrease of HA *Ab* “unmasked” the impact of CA *Ab* isolates. Similar “unmasking” events may explain other reports of increased proportions of *Ab* infections occurring outside of hospitals over time [24]. A limitation of our ecological study design is that we did not determine whether isolates were associated with clinical disease or asymptomatic colonization, so the impact of this CA reservoir on *Ab* disease remains to be determined. Regardless, as aggressive measures against nosocomial infections are implemented, future investigations should differentiate between outpatient *Ab* reservoirs, a microbial population neglected by investigations that largely focus on HA *Ab* infections, and “classical” nosocomial *Ab*.

Our comparative analysis begins to define the *Ab* community reservoir. Though there were no differences in patient age or sex between CA and HA cases (**Table 1**), we observed that CA isolates were most often from SST/MSK or urinary sources and that HA isolates were predominately from respiratory sources (**Figure 6B**). This is consistent with observations from a Hong Kong teaching hospital, where 32.8% and 25.8% of general ward *Acinetobacter* isolates were from wound or urinary sources, while 80.7% of ICU isolates were respiratory [25]. Similar observations were made in *Ab* populations in Spanish hospitals [24]. Though the anatomic source of isolates are probably influenced by the variable culturing practices inherent to different hospital wards, the fact that various international studies reported similar findings supports that these observations reflect real ecological phenomena.

In contrast to the susceptible *Ab* strains implicated in community-acquired pneumoniae in tropical regions [26], BJC CA *Ab* isolates displayed elevated CRAb/MDR rates (∼40%), albeit lower rates than HA isolates (∼60%) (**Table S3**). The high but differing resistance rates between BJC CA and HA *Ab* are consistent with rates reported in prior U.S. national studies [16, 18]. However, antibiotic pressures alone may not explain the diverging epidemiology of HA and CA *Ab*, as associations between anatomic source and CA or HA *Ab* were mostly conserved across CRAb and non-CRAb isolates (**Figure 6B**). It has been proposed that *Ab* capable of human colonization compose clonal subsets distinct from *Ab* occupying undefined environmental reservoirs [27]. It is tempting to speculate that clinically-relevant *Ab* subpopulations exhibit diverging capabilities to survive in different epidemiologic compartments or host niches, independent of antibiotic resistance. Examining this hypothesis will require molecular and phenotypic analyses of *Ab* isolated from different epidemiological compartments.

In summary, we report divergent antibiotic transmission dynamics, anatomic sources, and resistance rates between clinically-relevant CA and HA *Ab* populations. Though our findings are limited to a single regional U.S. healthcare system, similar observations have been reported by multiple groups nationally and internationally. Validating a cutoff of 48 hours post-hospital admission to define HA *Ab* subgroups requires more comprehensive review of *Ab* clinical cases (e.g., identifying HCA cases among CA isolates, clinical outcome analyses, etc.) coupled with molecular characterization of matched isolates. As endemic, non-nosocomial MDR *Ab* reservoirs pose potential threats to ongoing efforts against MDR *Ab* disease, further characterization of CA *Ab* isolates remains crucial.

## Acknowledgments

The authors would like to thank Dorothy Sinclair and Cherie Hill for their essential and expert contributions in data retrieval for this study.

## Abbreviations

Ab: *cinetobacter baumannii*
BJC: BJC HealthCare System
BJH: Barnes-Jewish Hospital
CA: community-acquired
CDR: Clinical Data Repository
CRAb: carbapenem-resistant *A. baumannii*
HA: hospital-acquired
HCA: healthcare-associated
MDR: multidrug-resistant
SST/MSK: skin and soft tissue/musculoskeletal

---

**Figure S1.**
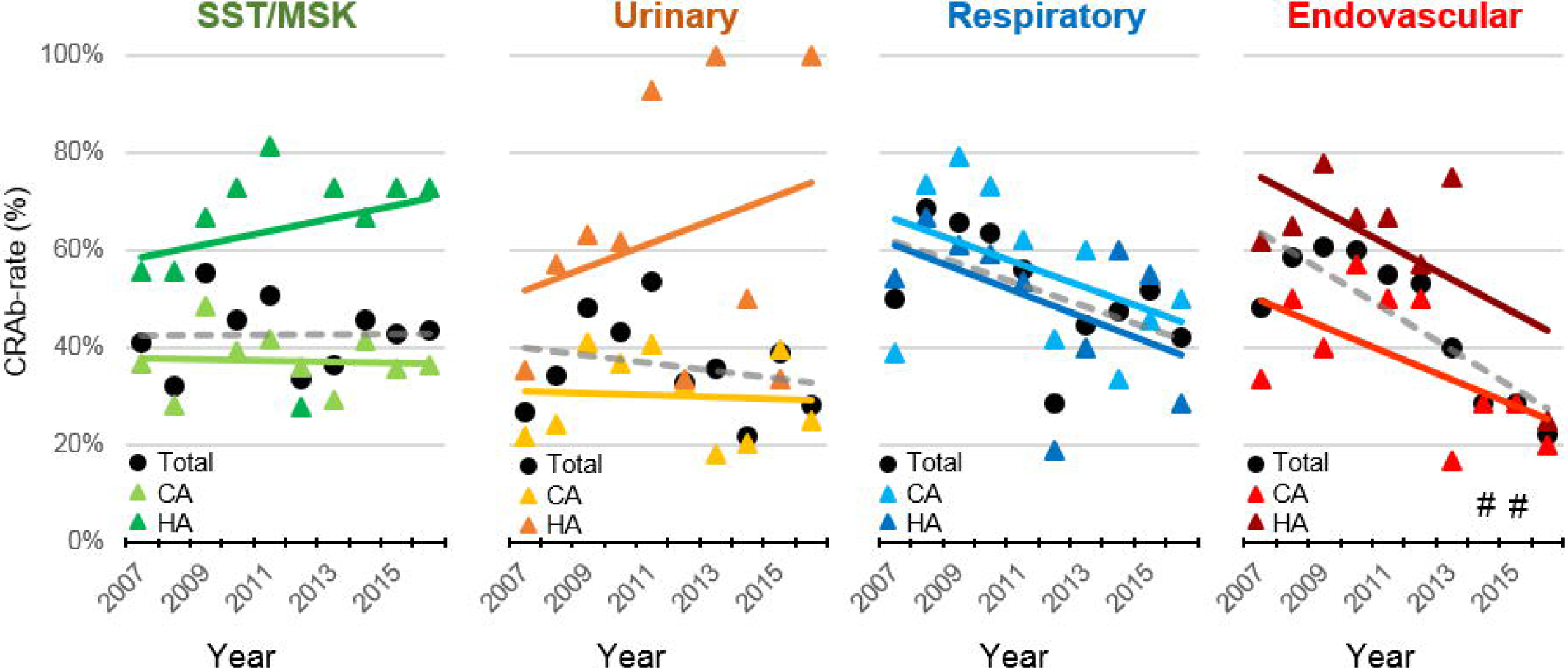
Rates of carbapenem resistance (CRAb-rate) among *Ab* isolates from different anatomic sources, BJC 2007-2016. Annual CRAb-rates among total (black circles, gray dashed line), hospital-acquired (HA, darker triangles and solid line), and community-acquired (CA, lighter triangles and solid lines) *Ab* isolates, grouped by anatomic source. #, no HA endovascular isolates with carbapenem susceptibility data were identified in 2014 or 2015.

**Figure S2.**
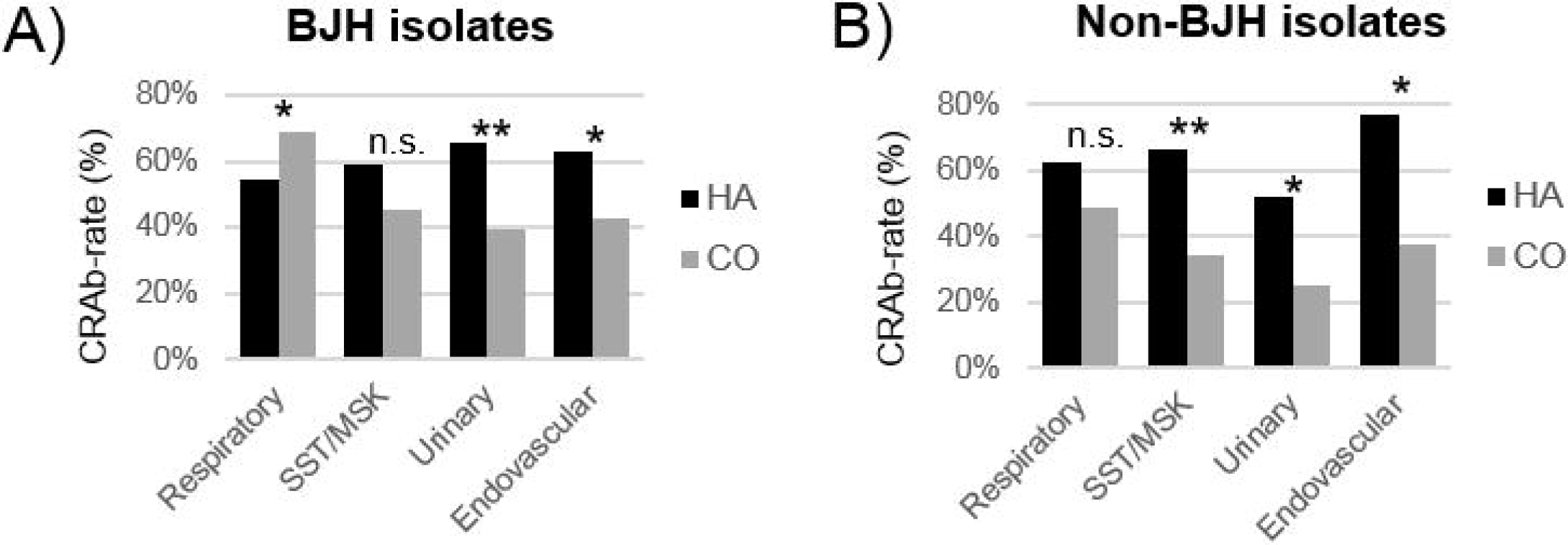
Rates of carbapenem resistance (CRAb-rate) among adult BJH (panel A) and non-BJH (panel B) isolates. CRAb-rates were compared between hospital-acquired (HA, black bars) and community-acquired (CA, gray bars) isolates from each anatomic source by chi-squared test. SST/MSK, skin and soft tissue/musculoskeletal; n.s., not significant; *, p<0.05; **, *p*<0.005. There was no CRAb-rate difference among HA isolates in either panel A or B (*p*=0.28 and 0.402, respectively).

**Figure S3.**
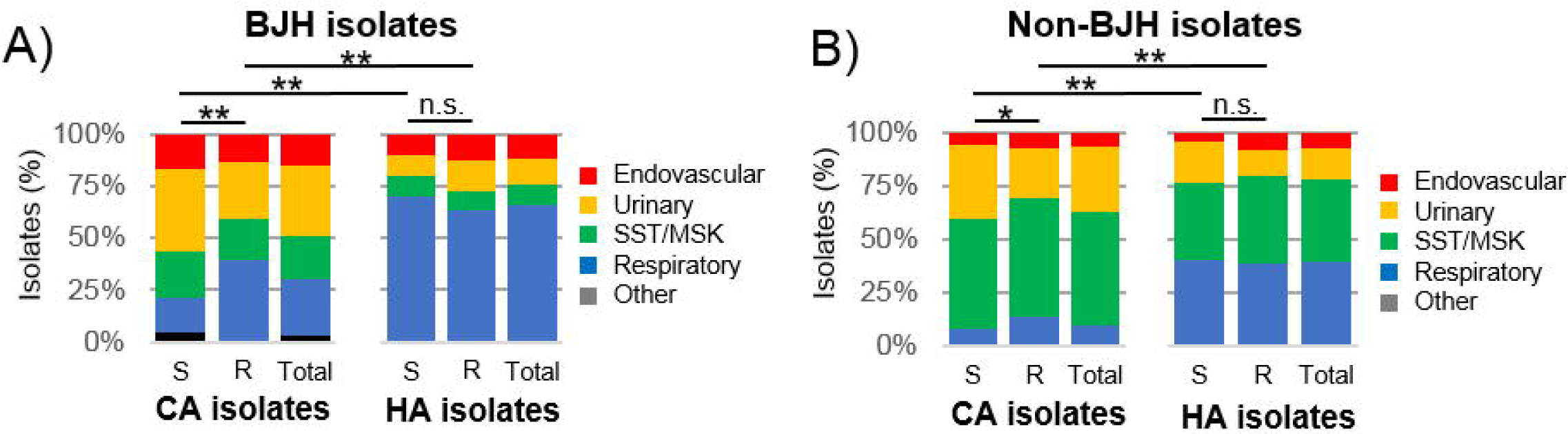
Proportion of carbapenem-susceptible (S), -resistant (R) or total adult isolates from each anatomic source, among BJH (panel A) and non-BJH (panel B) isolates. Graphs depict distributions among hospital-acquired (HA) and community-acquired (CA) isolates. The proportion of isolates from each source was compared between compartments by chi-squared test. SST/MSK, skin and soft tissue/musculoskeletal; n.s., not significant; *, p<0.05; **, *p*<0.005.

**Table S1.**
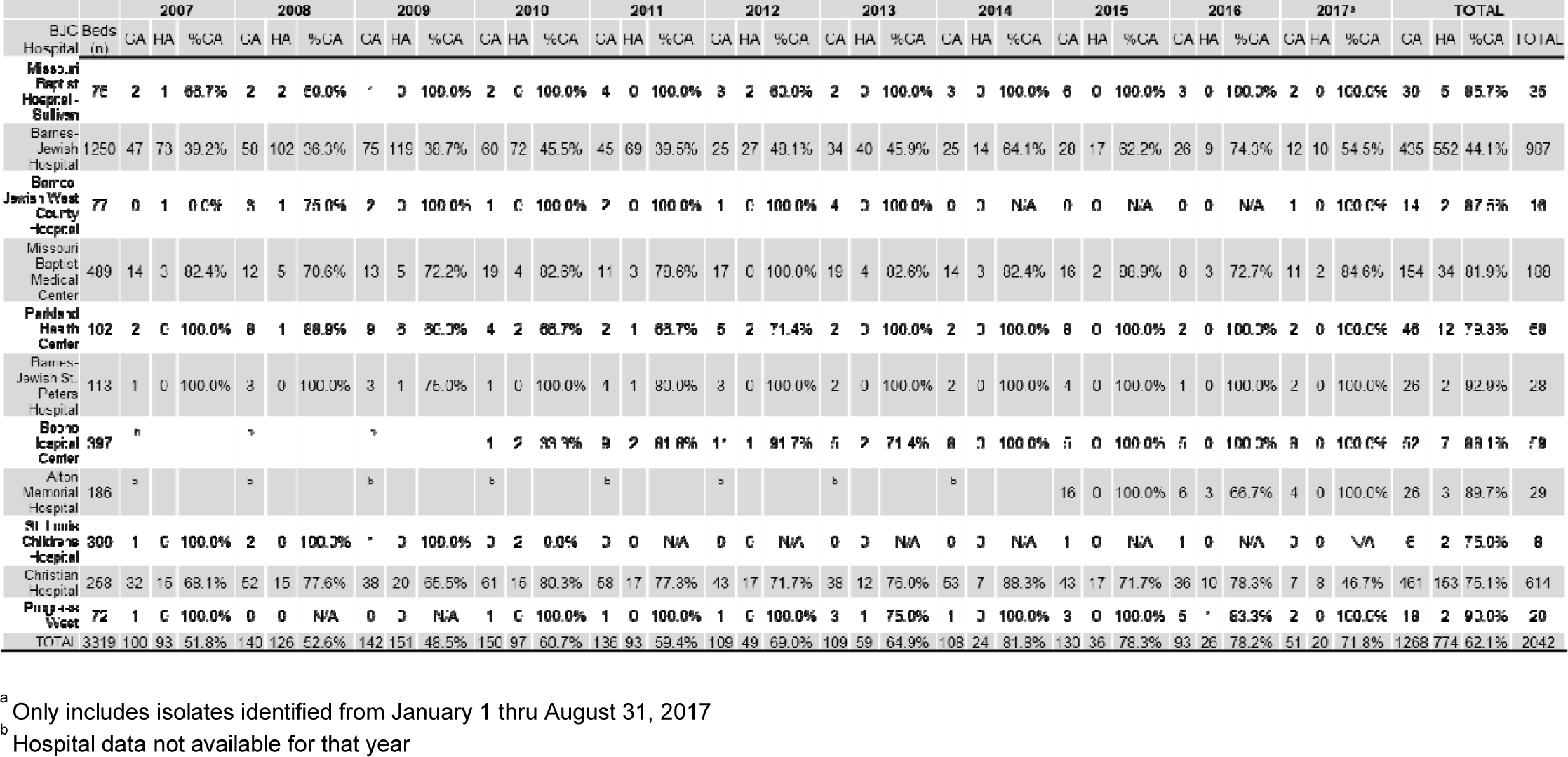
Annual amounts of community-acquired (CA) and hospital-acquired (HA) *A. baumannii* isolates per Hospital, BJC 2007-2017.

**Table S2.**
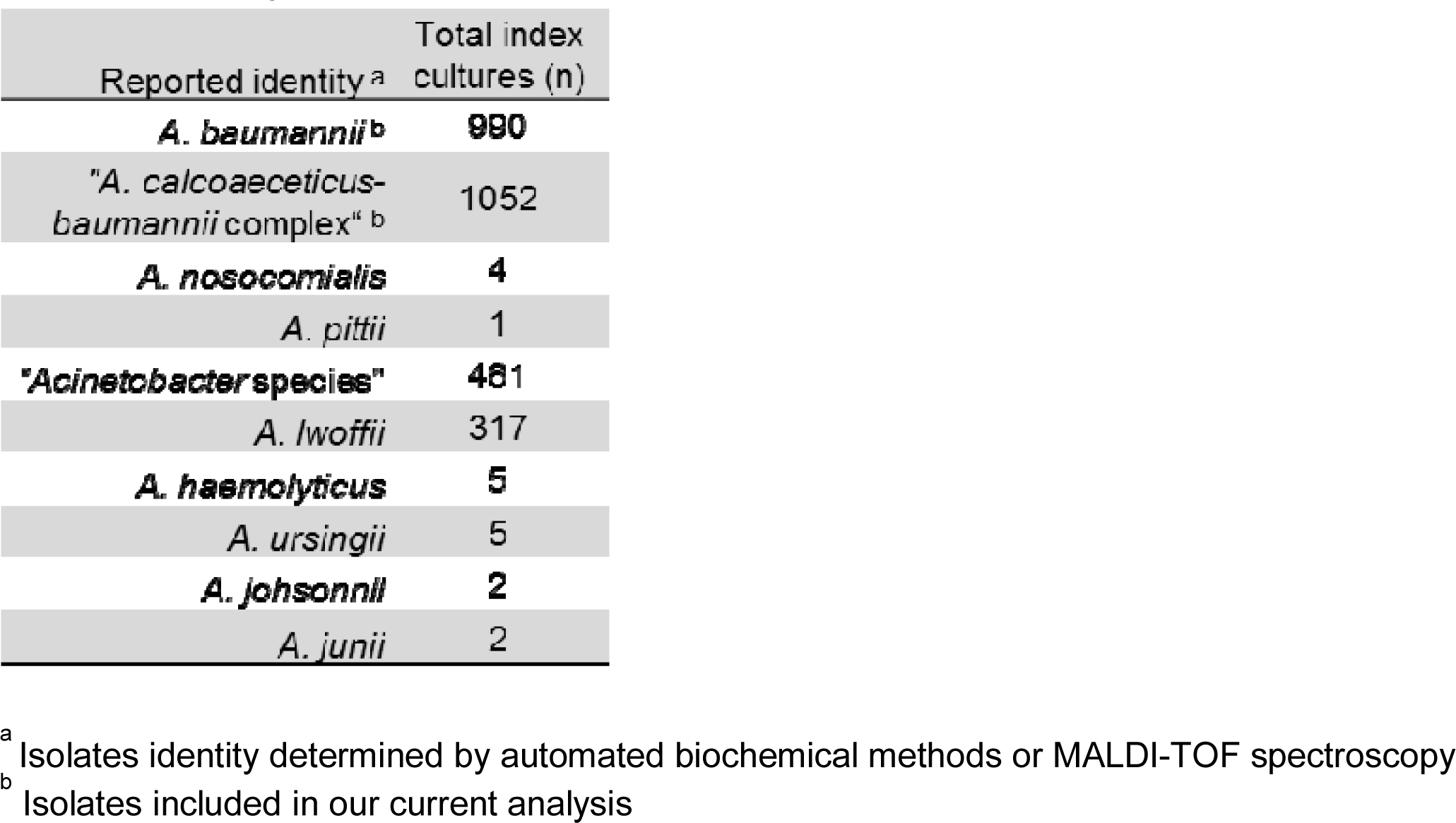
Identity and total amount of *Acinetobacter* index cases, BJC 2007-2017.

| Reported identity <sup>a</sup> | Total index cultures (n) |
| --- | --- |
| ***A. baumannii*<sup>b</sup>** | **990** |
| "*A. calcoaceticus*-*baumannii* complex"<sup>b</sup> | 1052 |
| ***A. nosocomialis*** | **4** |
| *A. pittii* | 1 |
| **"*Acinetobacter* species"** | **481** |
| *A. lwoffii* | 317 |
| ***A. haemolyticus*** | **5** |
| *A. ursingii* | 5 |
| ***A. johnsonii*** | **2** |
| *A. junii* | 2 |

<sup>a</sup> Isolates identity determined by automated biochemical methods or MALDI-TOF spectroscopy
<sup>b</sup> Isolates included in our current analysis

**Table S3.**
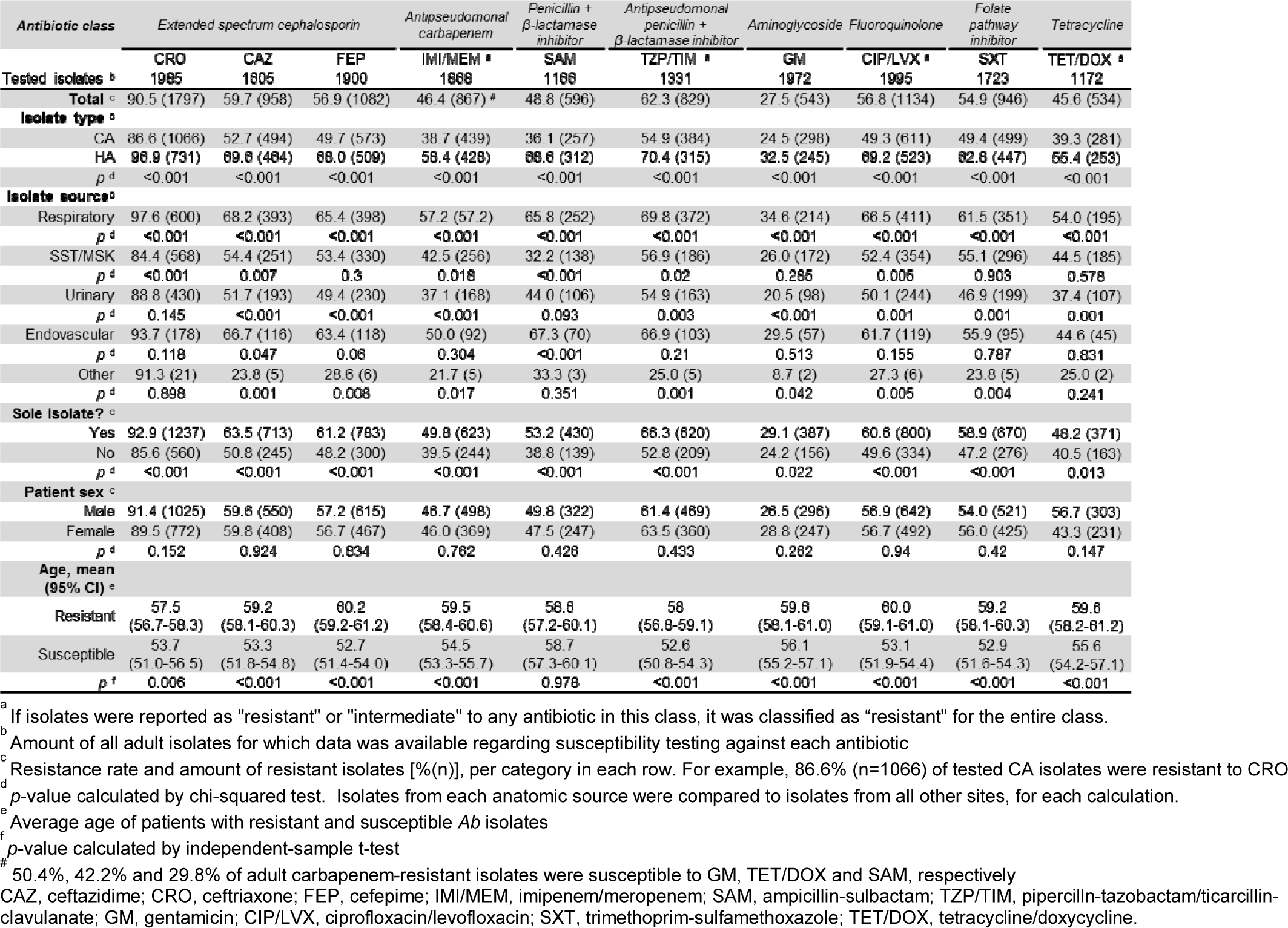
Associations between clinical characteristics and antibiotic resistance among all adult *A. baumannii* isolates, BJC 2007-2017.

